# Loss of PAX6 alters the excitatory/inhibitory neuronal ratio in human cerebral organoids

**DOI:** 10.1101/2023.07.31.551262

**Authors:** Wai Kit Chan, Shibla Abdulla, Lusi Zhao, Danilo Negro, Victoria M. Munro, Helen Marshall, Zrinko Kozić, Megan Brown, Michela Barbato, Mariana Beltran, Neil C Henderson, David J. Price, John O. Mason

## Abstract

The transcription factor PAX6 is a crucial regulator of multiple aspects of embryonic forebrain development. It has well-established roles in the regulation of excitatory and inhibitory neuron development in the embryonic cortex in mice but PAX6’s roles during human forebrain development are less well understood. Using human cerebral organoids, we investigated PAX6’s roles in human neurodevelopment. Homozygous PAX6 mutant (*PAX6^-/-^*) organoids were larger than controls and contained increased inhibitory cell types. Excitatory neurons were still generated in *PAX6^-/-^* organoids but they were less mature, and a subset showed dysregulated expression of inhibitory identity genes compared to *PAX6^+/+^* controls. The inhibitory cells found in *PAX6^-/-^* organoids physically segregated from excitatory neurons and presented a distinct transcriptomic profile when compared to *in vivo* cortical inhibitory neurons. *PAX6^-/-^* organoids showed a dysregulated cellular response to PTN-PTPRZ1 signalling, which contributed to the observed increase in inhibitory neurons and the consequent altered excitatory to inhibitory neuronal ratio.

**Summary Statement:** Human cerebral organoids lacking PAX6 expression show an altered excitatory to inhibitory neuronal ratio, concomitant with altered responses to intercellular PTN-PTPRZ1 signalling.

## Introduction

Neurons in the cerebral cortex fall into two major classes – excitatory (which express the neurotransmitter glutamate) and inhibitory (which express the neurotransmitter GABA) (Cadwell et al., 2019). Excitatory neurons are born in the dorsal telencephalon while large numbers of inhibitory neurons are born ventrally, in the ganglionic eminences before migrating into the cortical plate to form cortical circuits with excitatory neurons. (Anderson et al., 1997; Anderson et al., 2001; Gorski et al., 2002) Recent evidence suggests that in humans, some cortical inhibitory neurons are born in the cortex alongside excitatory neurons (Chung et al., 2024; Delgado et al., 2022; Keefe et al., 2025; Wang et al., 2025a). Imbalances between these two cell types are thought to underlie a variety of neurodevelopmental disorders (Bruining et al., 2020). The transcription factor PAX6 is a key regulator of embryonic forebrain development playing a crucial role in cell fate determination (Manuel et al., 2015; Ochi et al., 2022; Walcher et al., 2013). In humans, homozygous PAX6 mutations are either lethal or result in major brain defects including corpus callosum agenesis, cortical thinning, microcephaly, large midline cavities and hypoplastic brainstem (Glaser et al., 1994; Schmidt-Sidor et al., 2009; Solomon et al., 2009). Human PAX6 is predominantly expressed in dorsal regions of the ventricular zone (VZ) in the telencephalon, the region that gives rise to excitatory neurons (Kerwin et al., 2010; Lindsay et al., 2005). Transcriptomic and *in situ* hybridisation analysis of the human fetal cortex revealed high *PAX6* expression in the anterior areas of the cortex that decreased caudally (Eze et al., 2021; Kerwin et al., 2010). In the dorsoventral axis, PAX6 is highly expressed dorsally with a decrease ventrally towards the ganglionic eminences forming a dorsoventral gradient of expression. Human PAX6 is expressed at low levels within the striatal subpallium, partly colocalizing with the ventral transcription factor OLIG2 (Kerwin et al., 2010; Mo and Zecevic, 2008). At more caudal levels however, the pallial/subpallial boundary (PSPB) is clearly delineated by a sharp border between the PAX6-expressing pallium located dorsally, which gives rise to glutamatergic neurons, and the OLIG2-expressing subpallium located ventrally from which GABAergic neurons arise (Kerwin et al., 2010). Besides the VZ, PAX6 is also expressed in the subventricular zone (SVZ) in human, including both the inner and outer subventricular zones (iSVZ and oSVZ) (Eze et al., 2021; Kerwin et al., 2010; Nowakowski et al., 2017; Pollen et al., 2015). The oSVZ is populated by outer radial glia (oRG) cells that characteristically marked by high expression of PTPRZ1, a receptor for PTN (Hansen et al., 2010; Pollen et al., 2015; Reillo et al., 2011). However, the functional significance of PTN-PTPRZ1 signalling in oRGs during development is unknown. The expansion of oRGs, which express PAX6, is a key driver of cortical expansion and folding in human brain development due to their remarkable self-renewing capacity (Pollen et al., 2015). While the expression pattern of PAX6 in the developing human brain is reasonably well described, little is yet known about its functions in the developing human cortex. Studying cerebral organoids derived from *PAX6^-/-^* induced pluripotent stem cells (iPSCs), we found that PAX6 loss resulted in the generation of abnormal inhibitory cell types that segregate from other neuronal cell types and are transcriptomically abnormal. While excitatory neurons were still generated in the absence of functional PAX6, their maturation was delayed, and a subset of excitatory neurons were also transcriptionally abnormal. Thus, PAX6 loss resulted in organoids that have an altered excitatory to inhibitory ratio and we show that this was partially due to dysregulation of the PTN-PTPRZ1 signalling pathway.

## Results

### Generation of PAX6^-/-^ iPSCs and organoids

PAX6 contains two DNA-binding domains, the paired box and homeodomain (Xu et al., 1999). Paired box mutations have been implicated in the brain malformations described in human patients (Tzoulaki et al., 2005) and the paired domain alone was found to be necessary and sufficient to regulate telencephalic development in mice (Haubst et al., 2004). Therefore, to study the role of PAX6 in human telencephalic development, we targeted the PAX6 loss of function mutation to the paired domain. We derived *PAX6^-/-^* mutant iPSC lines from the parental line NAS2 (Devine et al., 2011) using CRISPR-Cas9 to insert a T residue into the paired box region of PAX6 (Fig 1A), causing a frameshift mutation that results in premature termination of PAX6 protein at amino acid residue 31, 20 residues downstream of the mutation. Two independent *PAX6^-/-^* clones (A10 and B2) homozygous for this mutation and two isogenic control clones (Nas2 and Cas92) were selected for further analysis. We confirmed that PAX6 mutagenesis was successful by PAX6 immunostaining with an antibody specific for the paired box domain (Engelkamp et al., 1999) and western blot. The former confirmed absence of PAX6 paired domain in mutant organoids and the latter showed truncated PAX6 protein (Fig 1C-D). All iPSC lines used in this study were subjected to whole genome sequencing (WGS) analysis. No changes were detected at any of the five predicted most likely CRISPR off-target sites. Probing further to ascertain whether the targeting procedure may have introduced mutations that could have compromised the iPSC lines, we found that no impactful mutations that would affect pluripotency or genomic integrity of the iPSCs (Supplementary Fig 1). All lines expressed pluripotency markers NANOG, OCT3/4 and TRA-1- 60 (Supplementary Fig 2A)(Devine et al., 2011). Cerebral organoids were grown from *PAX6^+/+^* (control: NAS2 & CAS92) and *PAX6^-/-^* mutant (A10 & B2) cell lines using a version of the Lancaster protocol (Lancaster et al., 2013) modified by the inclusion of dual SMAD inhibition to enhance neural lineage differentiation (Chambers et al., 2009) and growth factors BDNF & NT3 to support growth and survival of neurons (Birey et al., 2017)(Supplementary Fig 2B).

**Fig 1.**
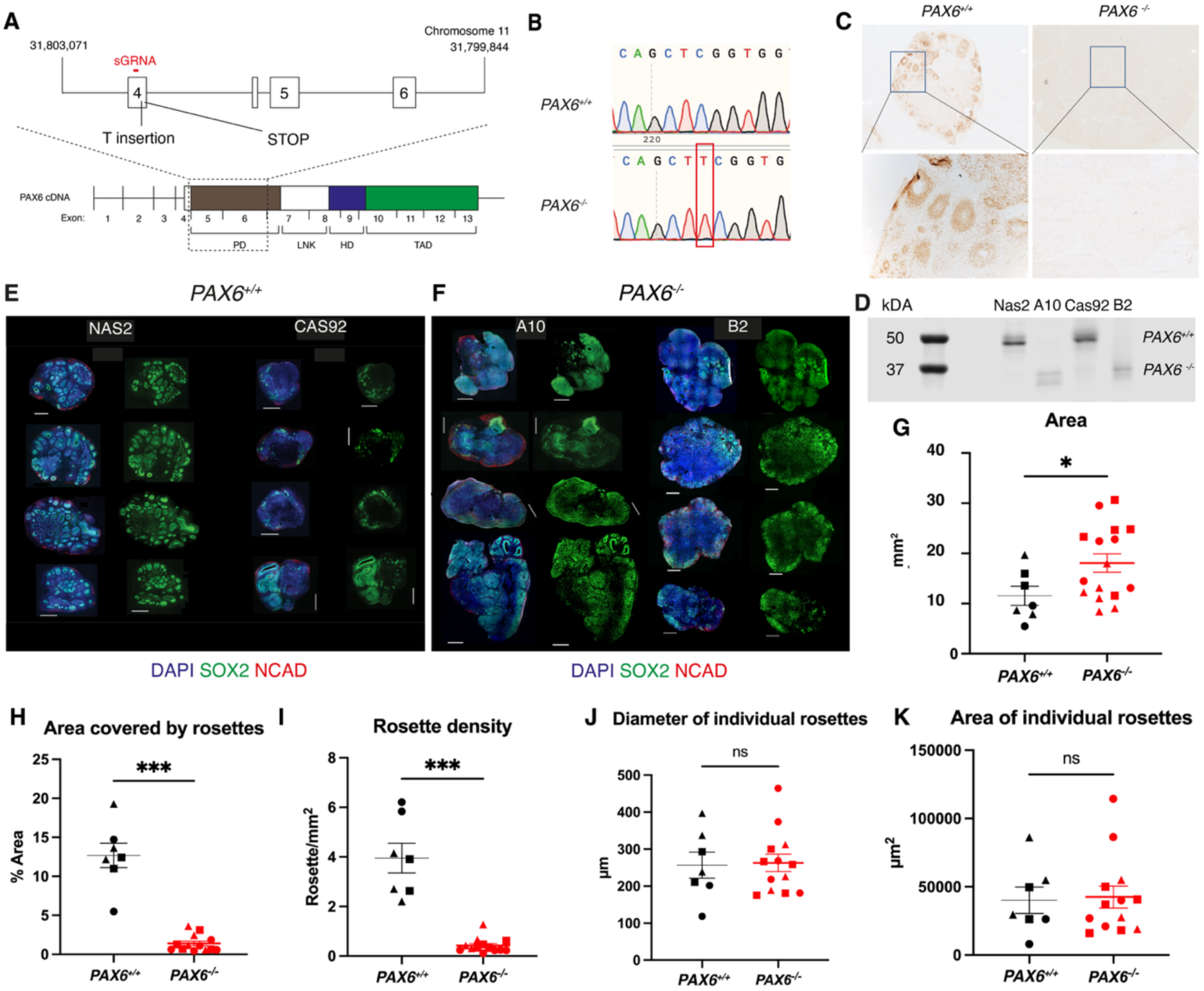
(A) Diagram showing part of the human *PAX6* locus indicating the location and nature of the mutation introduced to the *PAX6^-/-^* iPSC lines. Homozygous insertion of a single T residue in exon 4 caused a frameshift mutation leading to a premature stop codon within exon 4. Exons 1-3 are non-coding. PD – paired domain; LNK – linker; HD – homeodomain; TAD – transactivation domain. (B) Sanger sequencing of *PAX6^-/-^* iPSC genomic DNA showing the T insertion mutation. (C) PAX6 PD immunostaining of day 60 control and *PAX6^-/-^* organoids, confirming PAX6 PD domain absence in mutants. (D) Western blot showing PAX6 protein expression in *PAX6^-/-^* iPSCs, A10 and B2 and control lines Nas2 and Cas92. (E,F) SOX2 and N- CAD Immunostaining of representative sections of individual day 60 control and *PAX6^-/-^* organoids from separate differentiation batches (4). Scale bar = 1 mm. DAPI in blue, SOX2 in green, NCAD in red (G) Quantification of cross-sectional area of day 60 control and *PAX6^-/-^* organoids, p = 0.025 (H–K) Quantification of organoid size and rosette characteristics. (H) Area covered by rosettes between genotypes is significantly different with p = 0.0003. (I) Rosette density between genotypes was also significantly different with p = 0.001. (J – K) Neither the diameter or area of individual rosettes was significantly different between genotypes (p = 0.88 and p = 0.85 respectively). Each symbol represents an individual organoid while triangles, dots and square symbols indicate separate organoid batches. N ≥ 7 organoid batches. Each data point shown here consists of an average of 3 organoids per batch. * denotes significant difference of p < 0.05.

### PAX6^-/-^ organoids were larger than controls but contained fewer rosettes

We characterised the cerebral organoids generated from the iPSCs. To characterise the morphology of day 60 organoids grown from both *PAX6^-/-^* mutant lines and two isogenic control lines, we cryosectioned and stained for SOX2 (to mark neural progenitors) and N-CAD (to identify neuroepithelium) (Fig 1E,F). Staining patterns were consistent between organoids of the same genotype. Although *PAX6^-/-^* organoids appeared larger than controls, they showed a lower density of rosettes (radial arrangements of columnar epithelial cells with a central lumen resembling a cross-section of the developing neural tube). Organoid size was approximated by measuring the area of the widest cross-section of each organoid. The cross- sectional area of *PAX6^-/-^* organoids was 18.09 ± 1.80 mm^2^ compared to 11.55 ± 2.40 mm^2^ in controls (Fig 1G). The proportion of organoid tissue containing neural rosettes (demarcated by SOX2 and N-CAD staining (Gomes et al., 2020)) was 1.4% ± 0.29% in *PAX6^-/-^* organoids and 12.68% ± 1.57% in controls (Fig 1H). This decrease was due to a dramatically lower density of rosettes in *PAX6^-/-^* organoids (Fig 1I). While *PAX6^-/-^* organoids had lower neural rosette density, the size of individual rosettes did not differ significantly between *PAX6^-/-^* organoids and controls (Fig 1J-K). We previously reported that *Pax6^-/-^* mouse cortex showed an increase in inhibitory cell types that formed ectopias in the developing mouse cortex. These inhibitory cell types do not integrate into the cortical plate (Manuel et al., 2022). Therefore, we hypothesise that the increase in overall organoid size coupled with a decrease in neural rosette formation in *PAX6^-/-^* organoids was due to the emergence of ectopic inhibitory cell types. As organoids lack an anato mical axis (Cederquist et al., 2019), we define ectopic inhibitory cell types in organoids as inhibitory cell types whose distribution do not interleave with excitatory neurons as is usually seen in healthy cerebral organoids (Osaki et al., 2024; Sharf et al., 2022) but instead form separate territories in this study.

### Altered cellular composition in PAX6^-/-^ cerebral organoids

To characterise the cell types present in *PAX6^-/-^* and control organoids, we performed single cell RNA sequencing (scRNAseq) on four independent batches of organoids with each batch containing organoids grown from each of the four iPSC lines (two mutants: A10 and B2, and two controls: NAS2 and CAS92). Following quality control steps to remove ambient RNA, dead cells, doublets and stressed cells regularly found in brain organoids (Vértesy et al., 2022), 40,461 cells were retained for downstream analysis. Cells were annotated using CellTypist (Domínguez Conde et al., 2022) trained on a developing human fetal brain dataset (Braun et al., 2023). Annotation discrepancies were cross-checked against other models trained on human brain organoid data (He et al., 2024), other developing human fetal brain datasets (Wang et al., 2025a) and resolved using cell type specific marker genes described in previous studies (Bhaduri et al., 2020; Braun et al., 2023; He et al., 2024; Shi et al., 2021; Uzquiano et al., 2022; Yu et al., 2021) (Fig 2A,B). Our dataset contained radial glial progenitors (RGP) expressing marker genes including *GLI3*, *VIM* and *SLC1A3*; intermediate progenitor cells (IPC) expressing *NHLH1*, *HES6*, *EOMES*; two clusters of excitatory neurons: excitatory neuron 1 (EN1) expressing *NEUROD2, NEUROD6, SLC17A7* and excitatory neuron 2 (EN2) expressing marker genes including *NRXN1*, *RBFOX3*; inhibitory neuron progenitors that express *DLX2*, *DLX6, TOP2A* and inhibitory neurons that express *GAD1, SLC32A1,* and *DLX2* (Fig 2B, Supplementary Table 1). There were also two clusters containing small numbers of cells (<410). One of these (110 cells) was annotated as “unknown” as CellTypist was unable to reach agreement over the different trained models and none of the marker genes pointed towards a clear cell type (Supplementary Table 1). The other cluster (404 cells) was annotated as “mixed” because CellTypist annotated them as a mixture of excitatory and inhibitory neurons depending on the reference model. However, marker genes in that cluster included *DDIT4* and *HSP90AB1* which are typically found in stressed cells in organoids (Supplementary Table 1) (Vértesy et al., 2022). Both “mixed” and “unknown” clusters were found in both *PAX6^+/+^* and *PAX6^-/-^* organoids and were excluded from further analysis.

**Fig 2.**
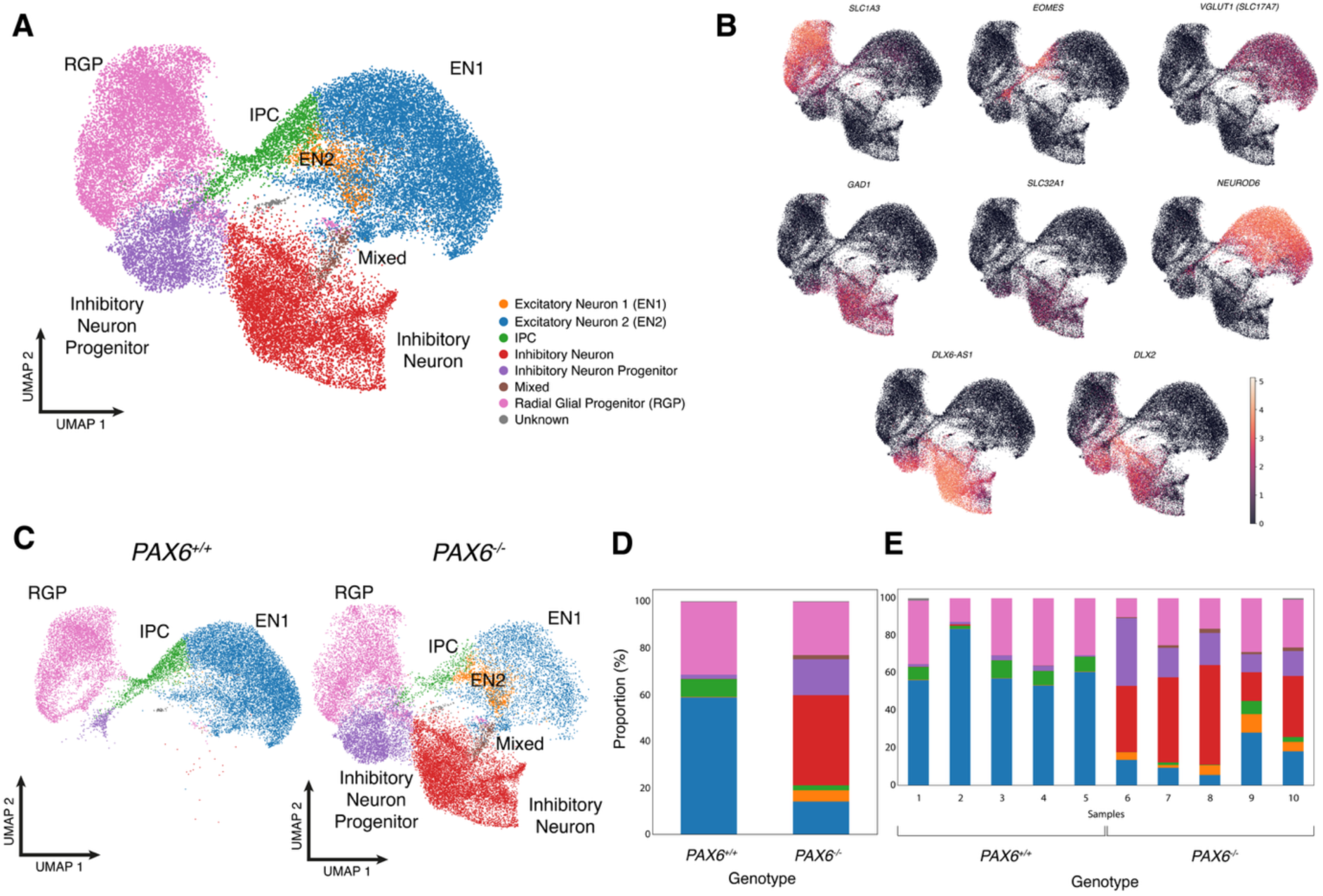
(A) UMAP plot showing combined scRNAseq data from day 60 *PAX6^-/-^* and control organoids. RGP – Radial glia progenitor; IPC – Intermediate progenitor cells; EN1 – excitatory neuron 1, EN2 – excitatory neuron 2. (B) UMAPs showing selected marker gene expression in specific cell types in day 60 organoids. (C) Separate UMAP plots of scRNAseq data from control (left) and *PAX6^-/-^* (right) organoids. (D-E) Comparison of cell type composition between control and *PAX6^-/-^* day 60 organoids showing all cell types coloured as in previous panels (A). (D) Comparison of cell types between *PAX6^+/+^* and *PAX6^-/-^* (E) shows cell compositions of individual samples

There was a change in the proportion of cell types found in PAX6-null organoids compared to controls (Fig 2C). The proportions of each cell type found in *PAX6^+/+^* organoids were: RGP 28.7 ± 3.7%; IPC, 6.7 ± 1.2%; EN1, 62.1 ± 4.9%; EN2 0.2% ± 0.01%; inhibitory neuron progenitors, 1.8 ± 0.3%; and inhibitory neurons 0.2% ± 0.1%.

In *PAX6^-/-^* organoids the proportions were: RGP 21.4 ± 3.1%; IPC, 2.3 ± 1.1%; EN1, 14.9 ± 3.5%; EN2, 5.1 ± 1.4%; inhibitory neuron progenitors, 18.4 ± 0.3%; and inhibitory neurons 36.5% ± 6.4%. Thus, *PAX6^-/-^* organoids showed a ∼25% decrease in RGPs and a ∼66% decrease in IPCs but a ∼922% increase in inhibitory neuron progenitors compared to *PAX6^+/+^* organoids (Fig 2C- E). There was also a ∼76% decrease in the number of EN1 neurons in mutant organoids (Fig 3C-E, blue). Inhibitory neurons (Fig 2C-E, red) were almost exclusively found in PAX*6^-/-^* organoids (36.5%) compared to *PAX6^+/+^* controls (0.2%), a ∼18150% increase in mutants. Finally, an excitatory neuronal cluster EN2 that was also almost exclusively found in P*AX6^-/-^* organoids, constituted 5.1 ± 1.2% of cells in PAX6-null organoids compared to 0.2 ± 0.01% in *PAX6^+/+^* control organoids (Fig 2C, EN2 - orange), a 2450% increase in mutants. Graphing the relative proportions of cell-types found in each individual sample showed that the cellular composition was relatively consistent within organoids of each genotype, as were the differences between *PAX6^-/-^* organoids and controls (Fig 2E).

**Fig 3.**
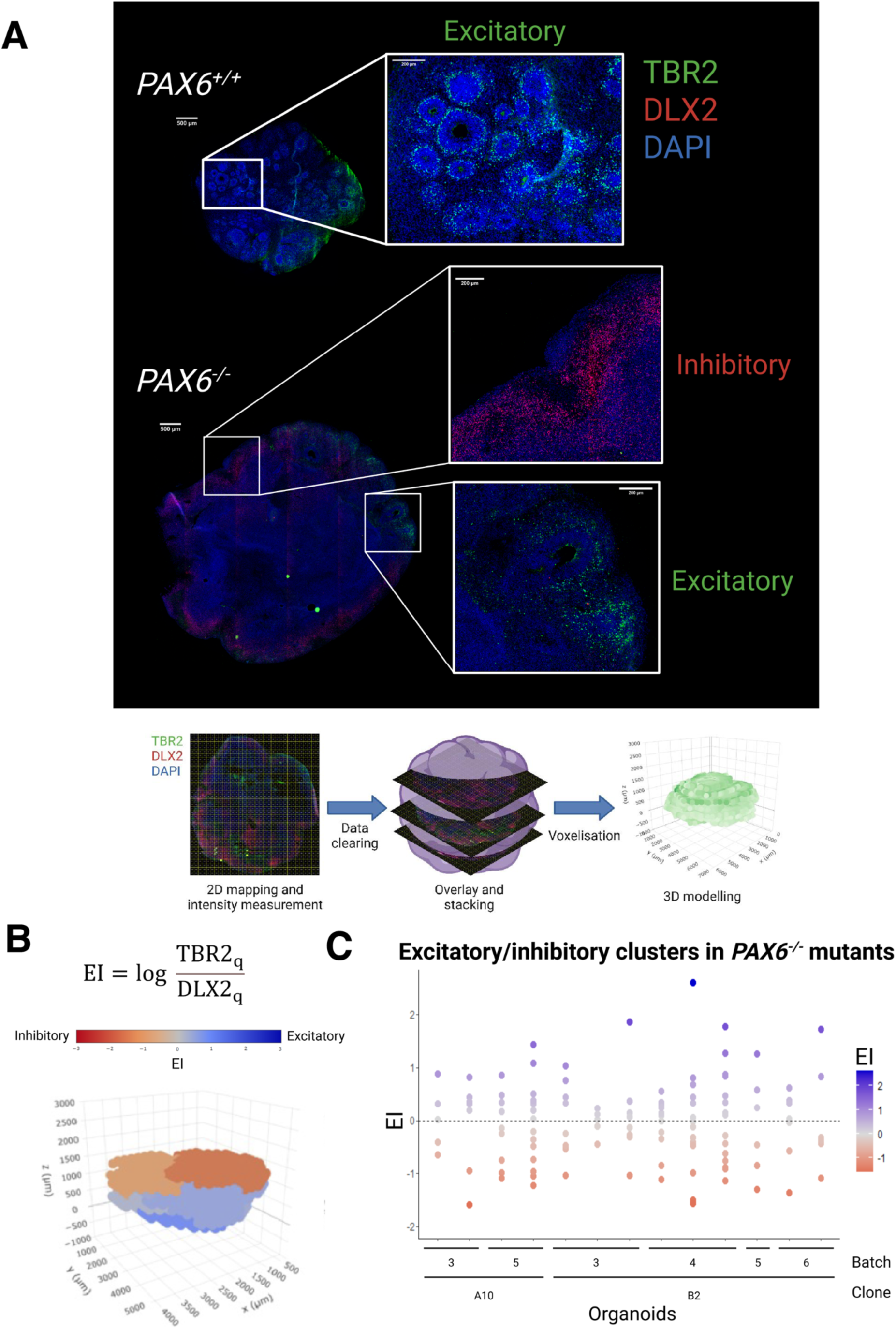
(A) Schema outlining method for production and quantitation of 3D models of organoids. Sections were immunostained for TBR2 and DLX2 and processed as shown. (B) Equation by which excitatory/inhibitory identities were conferred upon specific territories in organoids and (C) graph summarising the segregation of these territories in individual organoids.

In summary, we found that loss of PAX6 caused a reduction in the proportion of neural progenitors (RGP & IPC), a decrease in the proportion of excitatory neurons (EN1 & EN2) and an increase in the proportion of inhibitory progenitors and neurons (Fig 2D). The generation of inhibitory cell types at the expense of excitatory cell type suggests that PAX6 loss results in cell fate diversification of neural progenitors from an excitatory to an inhibitory fate.

### Inhibitory and excitatory cell types are physically segregated within PAX6^-/-^ organoids

While the scRNAseq results confirmed that loss of PAX6 results in the increased in inhibitory neurons, it did not address whether the inhibitory neurons were ectopic. To answer this question, we examined the distribution of excitatory and inhibitory identity cells in *PAX6^+/+^* and *PAX6^-/-^* organoids. We immunostained cryosections of day 60 organoids for TBR2 (EOMES) to label cortical IP cells which differentiate into excitatory neurons and DLX2 to identify inhibitory cell precursors (Fig 3A). Sections throughout each individual organoid were sampled and reconstructed to form 3D models (Fig 3A). Voxels in each reconstructed organoid were assigned a value corresponding to their relative level of expression of TBR2 or DLX2, giving a readout of dorsal versus ventral identity for cell clusters (Fig 3B, Supplementary Fig 3). We did not observe any inhibitory territory staining in the *PAX6^+/+^* organoids and our analysis clearly identified distinct, separate excitatory and inhibitory territories in each *PAX6^-/-^* organoid (Fig 3C). However, we did find some of these DLX2+ cells in the scRNAseq in the *PAX6^+/+^* control organoids, albeit a very small number (328 cells). This most likely indicates that detection of *DLX2* mRNA by scRNAseq is more sensitive than detection of DLX2 protein by immunofluorescence. Further, as we employed a sampling method of each organoid to build the 3D models, the small number of cells could be missed during sampling. Nevertheless, the separation of excitatory/inhibitory territories supports our hypothesis that PAX6 loss results in the emergence of ectopic inhibitory cell types. We next wondered whether PAX6 loss diverts neural progenitors only to specific inhibitory cell types.

### Inhibitory cell types in PAX6^-/-^ organoids are likely abnormal

To identify the specific subtype(s) of these ectopic inhibitory neurons that were generated in the *PAX6^-/-^* organoids, we integrated our dataset with a scRNAseq dataset from fetal human ventral telencephalon (Yu et al., 2021), where the majority of inhibitory neurons found in the cortex are generated, to determine whether the inhibitory cells found in the *PAX6^-/-^* organoids are equivalent to specific inhibitory subtypes found in the developing human brain (Fig 4A-B). For this analysis we focussed on the inhibitory neurons in the *PAX6^-/-^* organoids as they formed the vast majority of the inhibitory neurons in our dataset, 8704 cells compared to just 20 cells in the *PAX6^+/+^* control organoids. Co-clustering of inhibitory cells found in *PAX6^-/-^* organoids with a specific inhibitory subtype in the developing fetal brain would indicate transcriptomic similarity between them. However, we found that the inhibitory neurons in *PAX6^-/-^* organoids did not cluster with any specific subtype (Fig 4B). The failure of these cells to co-cluster with specific known fetal brain inhibitory subtypes suggests that they represent an aberrant population with a distinct transcriptomic profile, rather than mis-specified normal inhibitory neuron subtypes.

**Fig 4.**
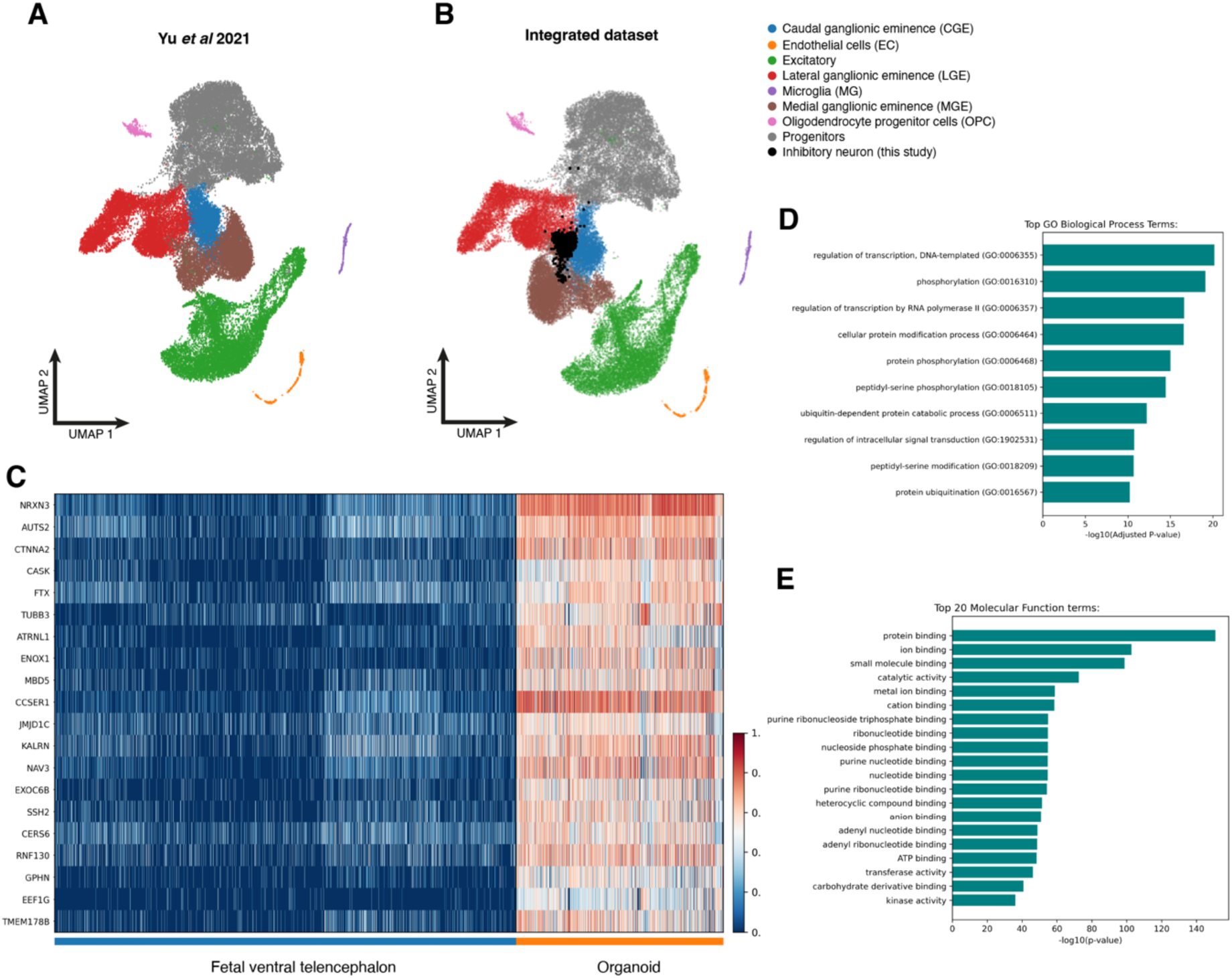
(A) scRNAseq dataset of fetal ganglionic eminences from (Yu et al., 2021). (B) Integrated dataset of inhibitory neurons (black) found in this study with the dataset from (A) showing that inhibitory neurons found in *PAX6^-/-^* organoids do not cluster to a specific inhibitory subtype found in *in vivo* fetal ganglionic eminences but a mixture of subtypes. (C) Heatmap showing top 20 DEGs between the inhibitory neurons from *PAX6^-/-^* organoid and fetal ventral telencephalon. (D) Top 10 biological process GO terms from the list of DEGs between *PAX6^-/-^* organoid and fetal inhibitory neurons. (E) Top 20 molecular function GO terms from the list of DEGs between *PAX6^-/-^* organoid and fetal inhibitory neurons.

To further explore the nature of *PAX6^-/-^* inhibitory cells in our *PAX6^-/-^* organoids, we conducted differential gene expression (DGE) analysis between *PAX6^-/-^* organoid inhibitory cells and inhibitory cells in the human ventral telencephalon reference dataset. This revealed a total of 6,933 differentially expressed genes (DEGs), 337 of which were downregulated in the *PAX6^-/-^* organoids compared to the reference dataset and 6,596 were upregulated (Supplementary Data). Plotting the top 20 DEGs in a heatmap revealed that all 20 were upregulated in *PAX6^-/-^* inhibitory cells (Fig 4C). Some DEGs, including *AUTS2*, *CASK* and *MBD5* are known to be involved in neuronal development while others encode cellular membrane proteins implicated in cellular signalling or cell morphology (*NRXN3*, *CTNNA2*, *TUBB3*, *ATRNL1* and *ENOX1*) (Fig 4C). To get a more holistic view of the differences between inhibitory cells in *PAX6^-/-^* organoids and those in the developing fetal brain, we performed gene ontology (GO) analysis on all 6933 DEGs. Top-ranked GO terms related to intra- and intercellular processes including transcription, phosphorylation and intracellular signal transduction (Fig 4D). Looking at the molecular function GO terms of the DEGs (Fig 4E), the main differences between inhibitory neurons generated from the loss of PAX6 and *in vivo* developing inhibitory neurons are dysregulated expression of genes involved in molecular interactions highlighted by most GO terms being involved in the binding of molecules.

Taken together, the DGE between *PAX6^-/-^* inhibitory neurons and in vivo inhibitory neurons suggests that the loss of PAX6 results in inhibitory neurons with defects in basic cellular processes including disrupted transcriptional regulation, cell signalling and molecular binding. The failure of *PAX6^-/-^* inhibitory neurons to establish proper molecular machinery for normal function suggests that the inhibitory neurons that were born due PAX6 loss were abnormal inhibitory neurons. Perhaps this might explain the extreme segregation into different “territories” as we have described above that usually happens to abnormal cell types (Fig 3).

### Excitatory cell types in PAX6^-/-^ organoids are more immature than those in PAX6^+/+^ controls

While inhibitory neurons numbers were increased in the expense of excitatory neurons in *PAX6^-/-^* organoids, the organoids still contained a significant number of excitatory neurons (Fig 3C-E). This indicates that loss of PAX6 diverted only a subset of progenitors to an inhibitory cell fate. We wondered whether PAX6 loss also affected the maturation trajectory of those excitatory neurons that are still being generated. To address this, we furthered our analysis into the excitatory neuronal clusters EN1 and EN2 (Fig 3). The majority of excitatory neurons in *PAX6^-/-^* organoids fell into cluster EN1 which was also found in controls (Fig 3C), indicating that these cells are transcriptionally similar to *PAX6^+/+^* cells. However, the number of EN2 cells were substantially increased (∼25 fold) in *PAX6^-/-^* organoids compared to controls (Fig 3C) suggesting that PAX6 promotes a subset of neural progenitors to acquire EN2 cell fate. How different are the cells in the EN2 cluster compared to EN1?

Highly expressed marker genes in EN2 cells include *NRXN1*, *CAMK4*, *DCC*, and *NDST3* which are all involved in cellular signalling. Further, key marker genes for excitatory neurons such as *NEUROD2* and *NEUROD6* (Bhaduri et al., 2020) are expressed at lower levels in EN2 cells than in EN1 (Supplementary Table 1, Supplementary Data). However, EN2 cells do express glutamate receptor genes including *GRID1*, *GRIK2* and *SHISA6* (Supplementary Table 1), characteristic of excitatory glutamatergic neurons. DGE analysis between EN2 and EN1 cells revealed higher levels of inhibitory marker genes including *DLX5* and *ASCL1*, and genes involved in cellular signalling including *NRXN1*, *PXDC1*, *ADAM22* (and many more) in EN2 cells (Fig 5A). Taken together, these findings indicate that a subset of excitatory neurons generated in *PAX6^-/-^* organoids have an atypical transcriptome, expressing both excitatory and inhibitory neuronal marker genes, perhaps as a consequence of dysregulated signalling and/or dysregulated cellular response to signalling.

**Fig 5.**
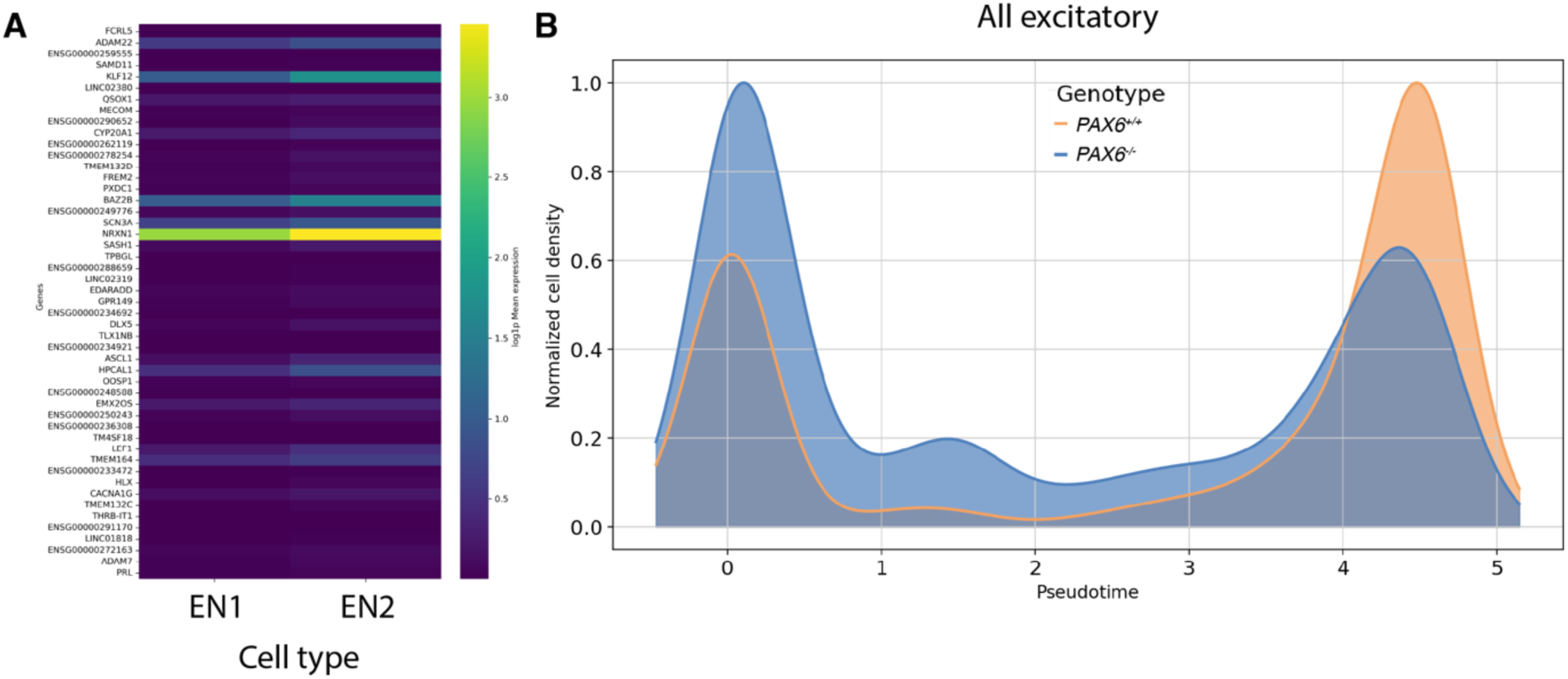
(A) Differential gene expression between EN1 and EN2 excitatory neuron clusters showing higher expression of inhibitory marker genes and genes involved in cellular signalling in EN2. (B) Pseudotime plot of excitatory lineage cells showing the developmental progression from progenitor to excitatory neurons. *PAX6^+/+^* is shown in orange and *PAX6^-/-^* is shown in blue, showing that *PAX6^-/-^* organoids have more immature cells along the excitatory lineage than do *PAX6^+/+^*organoids.

To determine whether PAX6 loss affects the lineage progression of excitatory neurons in organoids, we assessed the maturation of the excitatory lineage in both *PAX6^+/+^* and *PAX6^-/-^* organoids using pseudotime. EN1 and EN2 were grouped together as excitatory neurons for this analysis as there were insufficient EN2 cells in *PAX6^+/+^* organoids to allow a meaningful comparison. We found that *PAX6^+/+^* organoids contained more mature cells of the excitatory lineage than *PAX6^-/-^* organoids, as visualised by cell density along pseudotime (Fig 5B). Taken together, these results indicate that in addition to diverting a subset of cortical progenitors into an inhibitory lineage with an abnormal identity, loss of PAX6 results in a delay in the maturation of excitatory neurons and enrichment of a particular subset of excitatory neurons, EN2. This enrichment of EN2 however did not increase the proportion of total excitatory neurons in *PAX6^-/-^* organoids.

### Cell fate diversification in PAX6^-/-^ cerebral organoids may be driven by a dysregulated response to intercellular signalling

To address the mechanism underlying these cell fate diversifications caused by loss of PAX6, we identified genes which drive the diversification from excitatory to inhibitory cell fate in *PAX6^-/-^* organoids. We used scFates to infer developmental trajectories of RGPs (Faure et al., 2023) by computing pseudotime using RGP as a root node (Fig 6A-C). RGPs progressed towards either an excitatory (Fig 6D) or an inhibitory neuronal fate (Fig 6F). To identify driver genes in excitatory and inhibitory cell lineages, we performed DGE analysis along both developmental trajectories. Driver genes display dynamic, highly variable expression changes in the excitatory (Fig 6E) and inhibitory branches (Fig 6G). Plotting the expression of these genes along pseudotime for each branch, we found that they can be grouped into two categories: well-known marker genes for respective lineages and genes related to cellular signalling. For example, we identified marker genes for excitatory fate such as *NEUROD6* and *NEUROD2* (Fig 6E); and *DLX6-AS1*, *DLX5* and *ERBB4* for inhibitory fate (Fig 6G). Further, we found genes related to cellular signalling including *PTPRZ1*, *DLGAP1*, *THSD7A*, *DAB1* in the excitatory branch (Fig 6E); and *PDZRN3*, *SNTG1*, *NRXN3* in the inhibitory branch (Fig 6G). We chose to focus further analysis on the *PTPRZ1* pathway as this emerged as the top candidate driver gene in the excitatory trajectory (Fig 6E).

**Fig 6.**
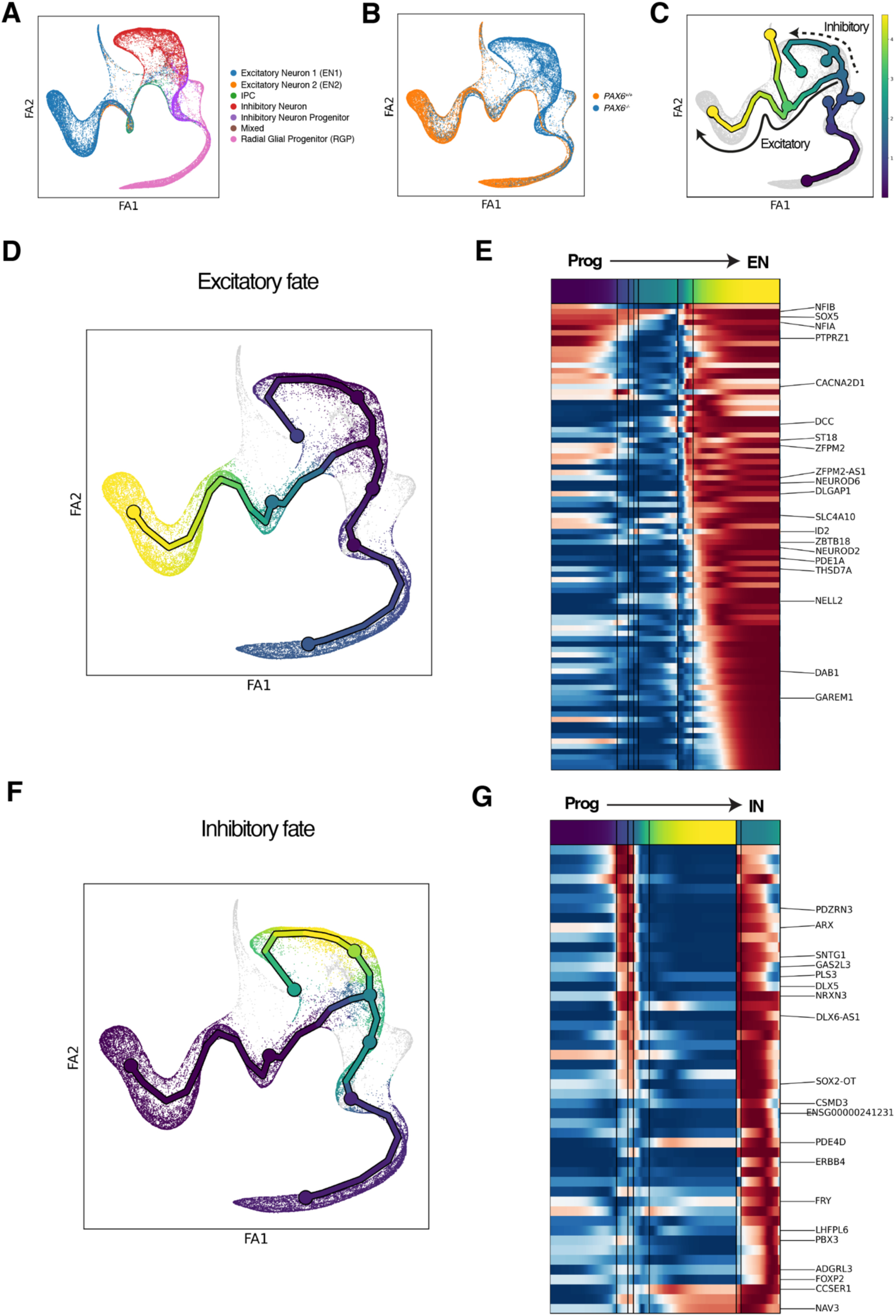
(A) Force-directed graph of scRNAseq data with cells arranged in pseudotime and coloured according to their cell type. (B) Same graph as in (A) coloured according to genotype, *PAX6^+/+^* in orange and *PAX6^-/-^* in blue. (C) Force-directed graph coloured in pseudotime with early pseudotime in dark blue and later pseudotime in yellow. (D) Force-directed graph showing excitatory differentiation trajectory and (F) inhibitory trajectory in pseudotime. (E) Expression pattern of genes from progenitor to excitatory neurons that were differentially expressed in the excitatory trajectory. (G) Expression pattern of genes from progenitor to inhibitory neurons that were differentially expressed in the inhibitory trajectory. Prog – progenitor; EN – excitatory neuron; IN – inhibitory neuron.

### Decreased sensitivity to PTN-PTPRZ1 signalling in PAX6^-/-^ cerebral organoids results in altered excitatory/inhibitory neuronal ratio

Protein tyrosine phosphatase receptor type Z1 (PTPRZ1) is a receptor for pleiotrophin (PTN) (Meng et al., 2000; Wang, 2020). PTPRZ1 is highly expressed in cortical progenitors, especially in oRGs, a progenitor cell type rarely found in mice but which is greatly expanded in primates (Pollen et al., 2015). PTN signalling has been implicated in neuronal development (Garcia- Gutierrez et al., 2014; Tang et al., 2019) and neurite outgrowth (Yanagisawa et al., 2010) but it has not previously been studied in the context of cortical development. To investigate whether the PTN-PTPRZ1 signalling pathway or the cellular response to PTN is dysregulated in *PAX6^-/-^* organoids, we first examined expression of *PTN* and its receptor *PTPRZ1* (Fig 7A-F). We saw no significant difference in *PTN* expression between *PAX6^+/+^* and *PAX6^-/-^* organoids (Fig 7A-C) but there was an overall decrease in *PTPRZ1* expression across all individual cell types in *PAX6^-/-^* organoids compared to controls (Fig 7D-F). In addition to PTPRZ1, PTN has been reported to interact with other cell surface receptors including ALK, SDC1, SDC2, SDC3, PTPRS, PTPRB, PLXNB2, CDH10 and NCL (González-Castillo et al., 2015; Wang, 2020). To be comprehensive, we also examined the mRNA expression of these receptors in our scRNAseq dataset, which revealed no significant differences between *PAX6^+/+^* and *PAX6^-/-^* organoids (Supplementary Fig 4).

**Fig 7.**
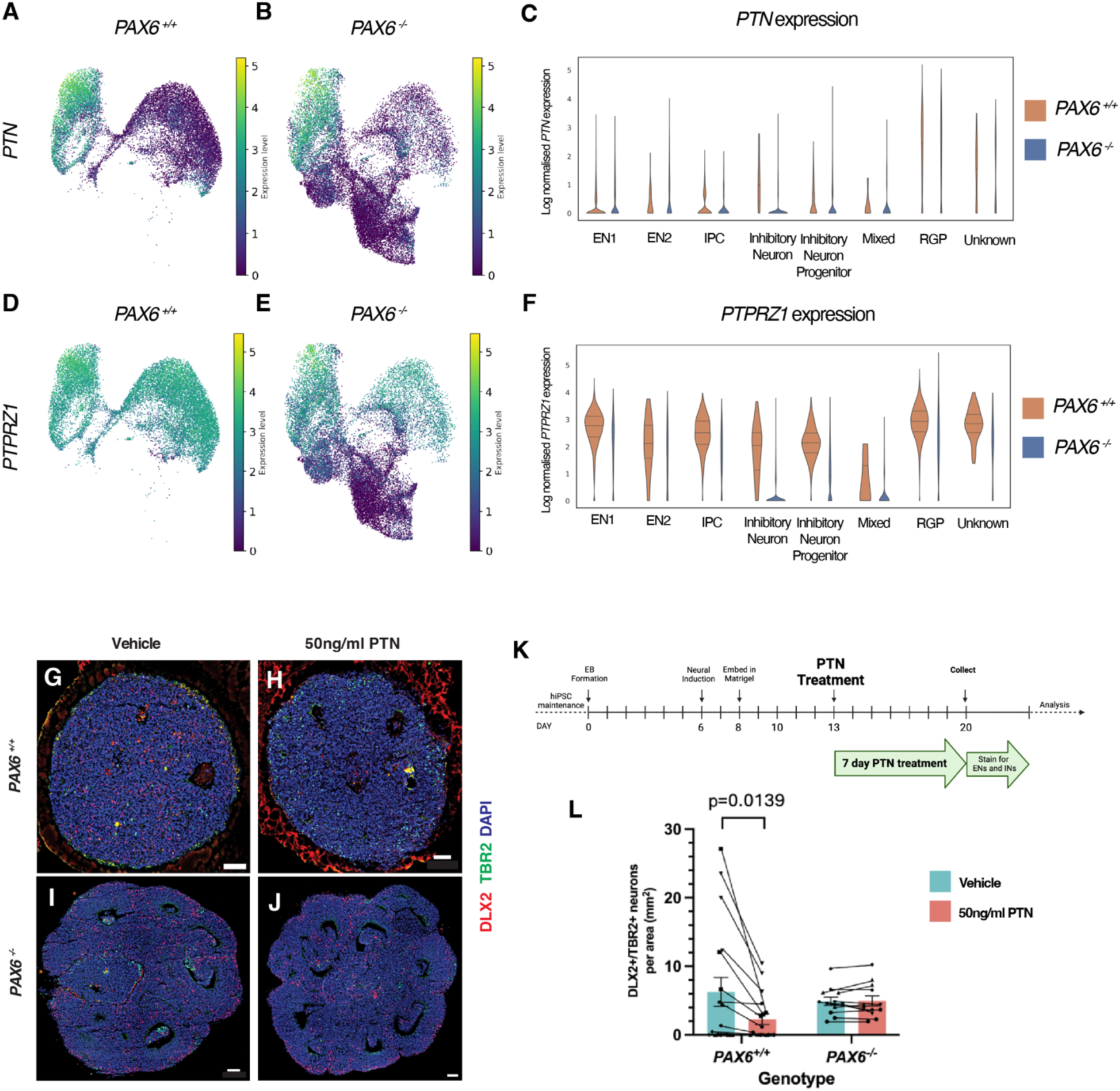
(A) UMAP plot of *PTN* expression in *PAX6^+/+^* and (B) in *PAX6^-/-^*. (C) Violin plots of *PTN* expression across different cell types found in each genotype, *PAX6^+/+^* in brown and *PAX6^-/-^* in blue. (D) UMAP plot of *PTPRZ1* expression in *PAX6^+/+^* and (B) in *PAX6^-/-^*. (C) Violin plots of *PTPRZ1* expression across different cell types found in each genotype, *PAX6^+/+^* in brown and *PAX6^-/-^* in blue. (G) Representative image of *PAX6^+/+^* organoid treated with vehicle and (H) treated with 50ng/ml PTN showing a decrease in DLX2+ cells (red). (I) Representative image of *PAX6^-/-^* organoid treated with vehicle and (J) treated with 50ng/ml PTN. (K) Schema showing PTN treatment timeline. (L) Quantification of DLX2+/TBR2+ cells per area after vehicle or PTN treatment.

We hypothesised that the lowered expression of *PTPRZ1* in *PAX6^-/-^* organoids could decrease their sensitivity to PTN signalling, leading to the misspecification of RGPs from excitatory to inhibitory fate. To test this possibility, we treated *PAX6^+/+^* and *PAX6^-/-^* organoids with PTN at an earlier developmental age (organoid day 20) that precedes our observations described above. After seven days, the ratio of inhibitory (DLX2+) to excitatory (TBR2+) cells was determined, as a readout of PTN-PTPRZ1 signalling sensitivity (Fig 7K). Treating *PAX6^+/+^* organoids with 50 ng/ml PTN (Fig 7G-H) resulted in a statistically significant 2.75-fold decrease (6.27 to 2.28) in the ratio of DLX2+/TBR2+ cells compared to vehicle treatment (Fig 7L). A surprising observation that we noted was the similarity of the DLX2/TBR2 before PTN treatment between D20 control and mutant organoids given that we described there were not many DLX2+ cells in D60 control organoids resulting in a stark difference in DLX2/TBR2 ratio between the control and *PAX6^-/-^* mutant D60 organoids (Fig 2-3). This difference likely reflects the developmental trajectory of cerebral organoids: early-stage organoids (<2 months) contain diverse cell types from multiple nervous system regions, including non-cortical cells, before transitioning to exclusively cortical cell types by ∼2 months/D60 (Uzquiano et al., 2022). This possibly explains why DLX2+ cells were present at D20 but drastically reduced at D60 in *PAX6^+/+^* control organoids resulting in a similar DLX2/TBR2 ratio between control and *PAX6^-/-^* mutant at D20 but not at D60.

Therefore, in the presence of PAX6, progenitor cells respond to the activation of the PTN- PTPRZ1 pathway by committing to an excitatory rather than inhibitory fate. However, treating *PAX6^-/-^* organoids with the same dose of PTN (Fig 7I-J) resulted in no change in the ratio of DLX2+/TBR2+ cells (Fig 7L) most probably due to lowered expression of PTPRZ1 rendering the cells less likely to respond to PTN. This indicates that dysregulated cellular response to PTN- PTPRZ1 signalling due to the loss of PAX6 contributes to the misspecification of RGPs from excitatory fate to inhibitory fate, thus altering the excitatory/inhibitory ratio.

## Discussion

Here, we have shown that PAX6 loss in human cerebral organoids leads to a subset of cortical progenitors deviating from excitatory to inhibitory fate, resulting in an altered excitatory to inhibitory neuronal ratio. The inhibitory cells generated from *PAX6^-/-^* physically segregate from excitatory cells, forming separate territories. While excitatory neurons were still generated in the absence of PAX6, their maturation appeared to be delayed and we found enrichment of transcriptomically atypical excitatory cells that show upregulated expression of inhibitory marker genes such as *ASCL1* and *DLX5* and genes important for cellular signalling such as *NRXN1*, *PXDC1*, *ADAM22*. Finally, we present evidence of dysregulated cellular response to the PTN-PTPRZ1 pathway in neural progenitors as a mechanism contributing to the altered excitatory to inhibitory neuronal ratio in *PAX6^-/-^* organoids.

### PAX6 mediated regulation of excitatory/inhibitory neuronal fate across model systems

Previously, we described Pax6’s role in ensuring that mouse cortical progenitors progress toward an excitatory neuronal fate by modulating the competence of the progenitors to respond to the signalling molecules Sonic Hedgehog (Shh), which promotes the adoption of inhibitory neuronal fate and Bone Morphogenetic Protein (BMP) which promotes excitatory neuronal fate without any detectable changes in the expression of the ligand or receptors for Shh or BMP. The result of this dysregulation in neural progenitors’ competence to respond to Shh and BMP was an increase of progenitors adopting inhibitory cell fate in the expense of excitatory cell fate (Manuel et al., 2022).

Our present study using human organoids showed that the effect of PAX6 loss on the excitatory/inhibitory neuronal ratio looks to be conserved across species. It also highlighted some possible differences between the two model systems. We showed that dysregulated neural progenitor response to the PTN-PTPRZ1 signalling pathway contributed to the altered excitatory/inhibitory neuronal ratio in *PAX6^-/-^* organoids, most likely due to reduced expression of the PTPRZ1 receptor. PTPRZ1 is normally highly expressed in human/primate oRGs, a progenitor population that is not readily found in mice and PTN has been reported to act as a trophic factor responsible for maintaining the oRG stem cell niche (Pollen et al., 2015). While we did not annotate oRGs separately from RGPs in our scRNAseq dataset, there are oRGs present in both *PAX6^+/+^* and *PAX6^-/-^* organoids as shown by the expression of the oRG markers *HOPX*, *PTPRZ1*, *FAM107A* (Pollen et al., 2015) (Supplementary Fig 5B). The data we obtained suggests that PTN-PTPRZ1 signalling could be a possible human-specific mechanism of action underlying the altered excitatory/inhibitory ratio in *PAX6^-/-^* organoids as *Ptprz1* was not found to be differentially expressed between wildtype and Pax6 cortex-specific knockout mice (Manuel et al., 2022).

### PTN-PTPRZ1 role in cell fate commitment

The neurotrophic factor PTN is highly expressed in neural stem cells in the adult hippocampus and facilitates the maturation of newborn neurons (Tang et al., 2019). Part of its role in maturation includes the stimulation of neurite outgrowth (Yanagisawa et al., 2010) however, it has not previously been implicated in cell fate specification - specifically neurons of excitatory or inhibitory fate as shown in this study. Cell cycle length can contribute to changes in cell fate, for example in the process of Yamanaka reprogramming in which somatic cells undergo an ultrafast cell cycle of about 8 hours/cycle (Chen et al., 2015). Therefore, PTN- PTPRZ1 signalling could affect neural cell fate specification by influencing the rate of proliferation or cell cycle time. PTN’s actions on proliferation and cell cycle are context dependent. PTN signalling is needed for cell cycle progression in glioblastoma cells as it can stimulate glioblastoma stem cells during tumour growth (Chang et al., 2006; Shi et al., 2017), promote proliferation of neural stem and progenitor cells in the hippocampus (Li et al., 2023). In other cell types such as bovine chondrocytes, PTN inhibits proliferation (Tapp et al., 1999). Further experimentation will be required to fully describe the molecular pathway by which PTN-PTPRZ1 signalling influences cortical excitatory/inhibitory neural fate specification. It is possible that changes in other signalling pathways contribute to the altered excitatory/inhibitory neuronal ratio in *PAX6^-/-^* organoids as cell fate commitment is a complex process and likely involves many other factors and cofactors. PTN-PTPRZ1 signalling is almost certainly only one piece of a complex puzzle. Other signalling-related genes that were differentially expressed in *PAX6^-/-^* cells differentiating along the inhibitory pathway included *PDZRN3*, *SNTG1* and *NRXN3* (Supplementary Fig 5A) though experimental validation of these findings is beyond the scope of this study.

### Atypical inhibitory and excitatory cells generated in PAX6^-/-^ organoids

We described inhibitory cells and a subset of excitatory cells, EN2, generated in *PAX6^-/-^* organoids showing atypical transcriptomic profile. Some gene expression differences between cells in organoids and cells in embryos are to be expected, given likely differences between the *in vitro* culture environment and fetal environment *in vivo*. However, given the dysregulation in *PAX6^-/-^* cells’ cellular response to PTN-PTPRZ1 signalling resulting in altered excitatory/inhibitory ratio of neurons, we consider that it is more likely that PAX6 loss resulted in the generation of neurons that express inhibitory identity genes in the case of inhibitory neurons and excitatory identity genes for the subset of excitatory neurons in the EN2 cluster but are abnormal in their ability to interact with intra and intercellular signalling pathways due to differences their molecular features compared to *in vivo* inhibitory or excitatory neurons respectively.

In the case of the inhibitory neurons, previous reports suggested that *Pax6* deletion in mice resulted in the generation of immature olfactory bulb INs that are marked by the expression of the genes *Sp8, Scgn, Calb2* and *Tshz1* (Guo et al., 2019; Kroll and O’Leary, 2005). However, our previous study on *Pax6* deletion in mouse cortical progenitors produce inhibitory neurons that showed a different response to *Gsx2* loss to normal dLGE cells, cells that would mature into olfactory bulb INs indicating that the inhibitory neurons in *Pax6^-/-^* mice are abnormal. Results from the current study supports the latter suggestion as while PAX6 deletion in organoids generated INs, they showed an atypical transcriptomic profile. We further assessed for olfactory bulb IN identity cells (cells that co-express *SP8* and *SCGN*; *SP8*, *CALB2* and *TSHZ1*) in our scRNAseq dataset but we did not find many, if any, cells at all (Supplementary Fig 5D-E). Further, the extreme segregation from other cell types in the organoid also supports the idea that these inhibitory cells are abnormal with an aberrant transcriptomic profile as normal inhibitory neurons integrate into the developing cerebral cortex *in vivo* and in organoid/assembloid models (Osaki et al., 2024; Sharf et al., 2022; Wang et al., 2025b).

## Material & Methods

### Cell culture and organoid generation

iPSCs were maintained at 37°C with 5% CO_2_ in StemMACS iPS-Brew XF (Miltenyi) on Laminin 521 (Biolamina) coated plates and passaged with 0.5 mM EDTA (Life Technologies). Organoids were differentiated using a modified Lancaster protocol (Lancaster et al., 2013) which included addition of 10 µM SB431542 (Tocris) and 100 nM LDN-193189 (APExBIO) for neural induction and addition of 20 ng/ml BDNF and NT3 (Peprotech) to the medium at D30 for two weeks to aid neuronal maturation (Supplementary Fig 2B). Briefly, 9000 cells were aggregated using ultra-low attachment 96-well plates into embryoid bodies. These embryoid bodies were then incubated in neural induction media containing dual SMAD inhibition as detailed above at day six. Neural induction media was refreshed for day seven and organoids were embedded in Matrigel on day eight. Organoids were transferred to an orbital shaker and maintained in 37°C with 5% CO_2_ incubators for 60 days. All media change were done every other day.

### CRISPR/Cas9 mutagenesis

iPSCs were transfected with sgRNA 3’ UCCCUUUCCUCAGGUCACAG 5’ (Synthego) via nucleofection using program CA137 on a 4D Nucleofactor X (Lonza) as per manufacturer’s instructions. Clones were obtained by single cell dissociation of the transfected pool followed by screening for PAX6 mutation by Sanger sequencing of the *PAX6* locus. Successfully edited clones were re-sequenced and pluripotency test were performed using STEMlight Pluripotency kit (Invitrogen) (Supplementary Fig 1). Likely sites for off-target effects were identified using Cas-OFFinder (http://www.rgenome.net/cas-offinder/) and annotated in whole genome sequencing (WGS) data of all the iPSC lines. WGS data of all cell lines are deposited in ENA (accession: PRJEB63603).

### Immunohistochemistry

12µm thick cryosections of organoids were permeabilized in 0.1% Triton/PBS before incubation with primary antibodies at 1/200 dilution using appropriate serum as blocking buffer without antigen retrieval step: SOX2 (rabbit, ab92494 Abcam), TBR2 (sheep, AF6166 Bio-Techne), DLX2 (mouse, sc-393879 Santa Cruz); with antigen retrieval step: PAX6 (mouse, AD2.38, described in Engelkamp et al., (1999)), NCAD (mouse, 610920 BD Biosciences). All primary antibodies (except PAX6) were detected using AlexaFluor secondary antibodies (Invitrogen) at 1/200 dilution. PAX6 was detected using Vectastain ABC kit (Vector Labs). StemLight Pluripotency Antibody Kit (Stem Cell Technologies) was used for pluripotency testing. All immunohistochemistry was performed on ≥ 3 sections of individual control and *PAX6^-/-^* organoids. Number of organoids of each genotype ≥ 3 were used from ≥ 3 differentiation batches.

### Western blot

Snap-frozen cells were homogenized in RIPA buffer [10 mM Tris-HCl, 140 mM NaCl, 1 mM EDTA, 1% Triton X-100, 0.1% sodium DOC, 0.1% SDS, and Pierce protease inhibitor tablet, EDTA-free (A32965, Thermo Fisher Scientific)] in a bead mill 24 homogenizer (Fisher Scientific). Genomic DNA was removed with a Benzonase digest (E1014, Merck) for 15 min at room temperature. Protein concentration was quantified using DC Bio-Rad II (5000112, Bio-Rad) in a microtiter plate using a FLUOstar Omega plate reader (BMG Labtech). Samples were diluted in one-third volume of 4× Laemmli buffer (0.5 M tris-HCl, 8% SDS, 0.01% bromophenol blue, 40% glycerol, and 10% beta-mercaptoethanol) and denatured at 95°C for 10 min before being storing at −80°C. Protein samples (20 μg per lane) were resolved and transferred to nitrocellulose membranes (Bio-Rad). Total protein was measured using Revert 700 total protein stain (926-11010, LI-COR), and membranes were blocked in Intercept (TBS) blocking buffer (927-60001, LI-COR) for 1 hour at room temperature. Rabbit anti-PAX6 primary antibody (901301; BioLegend; 1:500) was incubated at 4°C overnight. Secondary antibodies [IRDye 800CW donkey anti-rabbit IgG secondary antibody (926-32213, LI-COR); 1:10,000] were incubated for 1 hour at room temperature before data were captured on a LI-COR Odyssey Classic. Western blots were replicated multiple times using 3 to 6 biological replicates (organoids) per genotype. Data were analyzed using Image Studio Lite (LI-COR) and Microsoft Excel.

### 3D modelling

Every 16^th^ section of individual organoids labelled with TBR2 and DLX2 was sampled using Zeiss Axio Scan Z1 slide scanner. A 200 x 200 µm grid was superimposed on each section and fluorescent intensities corresponding to both DLX2 and TBR2 expression were measured for each square. Expression data was then voxelised by assigning three-dimensional coordinates to each grid square. Voxels were processed using the 3D modelling package Plotly (Sievert, 2020) to produce 3D heatmaps of TBR2 and DLX2 expression in each organoid. To determine whether excitatory and inhibitory cells were mixed or segregated, voxels of TBR2 and DLX2 expression were used to calculate a EI value, given by the formula: log TBR2/DLX2 intensities (Fig. 3B). This gave a value range to allow plotting ‘excitatory’ areas, which contain excitatory neurons and ‘inhibitory’ areas where inhibitory neurons are enriched.

### Organoid dissociation and library preparation for single cell RNA sequencing

Four organoids of each genotype from four separate differentiation batches were submitted for scRNAseq. Briefly, each organoid were cut into smaller pieces with a scalpel and incubated at 37°C in papain solution (Worthington) for 30 minutes (Velasco et al., 2019) before single cell resuspension for downstream processing using 10X Genomics Chromium Next GEM Single Cell 3ʹ Reagent Kits v3.1 as per manufacturer’s instruction. No organoids were pooled. Sequencing was performed using Ilumina NovaSeq.

### PTN treatment

Treatment on organoids were done according to (Notaras et al., 2022) with slight adjustments. Briefly, organoids were treated with 50 ng/ml recombinant human PTN, reconstituted in distilled water or water as control and added to differentiation media for 7 days starting from day 13. The media was changed on days 13 and 14, and then every other day until they were harvested on day 20. Number of organoids of each genotype ≥ 3 were used from ≥ 3 differentiation batches.

### Preprocessing of Single-Cell Transcriptomics Data

Raw sequencing reads were aligned to the human reference genome (GRCh38, version 2024- A) using Cell Ranger (v8.0.1, 10x Genomics), and cell-by-gene expression matrices were generated. Ambient RNA contamination was removed using CellBender (v0.3.0), and potential doublets were identified and excluded using scDblFinder (v1.18.0). Genes expressed in fewer than 10 cells were excluded from downstream analyses. Only high-quality cells were retained based on stringent quality control criteria: cells with mitochondrial RNA content below 10%, ribosomal RNA content below 15%, hemoglobin RNA content below 5%, and between 500 and 10,000 detected genes were included. To further refine the dataset, Median Absolute Deviation (MAD) filtering was applied to several quality metrics, including total Unique Molecular Identifier (UMI) counts, number of detected genes, mitochondrial and ribosomal RNA content, and the proportion of reads mapping to the top 20 most highly expressed genes. For each metric, the median and MAD were calculated across all cells, and cells exceeding five MADs from the median in any category were classified as outliers and removed. Stressed cells were also excluded using the Gruffi package. Additionally, cells not expressing FOXG1 were filtered out to focus our analysis on telencephalic cells.

### Integration and Annotation of Single-Cell Transcriptomics Data

Data integration and batch correction were performed using scVI (scvi-tools v1.3.1.post1), which generated a unified latent representation from the combined datasets. Cell type prediction was performed using CellTypist, incorporating both its built-in “developmental brain” model (Braun et al., 2023) and custom models built from published datasets (He et al., 2024; Uzquiano et al., 2022; Wang et al., 2025a). Predicted cell labels were validated by cross-referencing with known marker genes.

### Differential Expression Analysis

Pairwise comparisons between all cell type pairs were conducted using the Wilcoxon rank-sum test as implemented in Scanpy.

### Pseudo-time and Bifurcation Analysis

Pseudotime calculation, cell developmental trajectory and branching were analysed using scFates (Faure et al., 2023).

### Mouse Data Integration

Mouse single-cell data from (Manuel et al., 2022) were integrated with our dataset using scVI for cross-species comparison.

### Whole Genome Analysis

Sequencing reads were quality-checked with FastQC (v0.12.1) and aligned to the human reference genome (GRCh38) using BWA-MEM (v0.7.18). BAM files were sorted, duplicates marked, and base quality scores recalibrated using GATK tools (version 4.6.1.0). Variants were called per sample using GATK HaplotypeCaller in GVCF mode, followed by joint genotyping with CombineGVCFs and GenotypeGVCFs. Variant calls were filtered using GATK VariantFiltration and functionally annotated with the Ensembl Variant Effect Predictor (VEP, version 113.0).

## Statistical analysis

For each between-organoid comparison (*PAX6^+/+^* vs *PAX6^-/-^*), an ANOVA with nested mixed effects was performed, accounting for the random effect of batches and clones on variance in a hierarchical manner. Specifically, p-values and confidence intervals were calculated using a Type III Analysis of Variance with Satterthwaite’s method.

## Data availability

Raw sequences for both scRNAseq and WGS were deposited in ENA (PRJEB63603).

## Code availability

All code used for data analysis in this article is available at Github (https://github.com/chaned/pax6_hom)

## Acknowledgements

We thank Tilo Kunath for sharing the NAS2 hIPSC line. We are grateful to Spasimir Shishinyov, Alexander Markov, and Kanrayat Benjasupawan for assisting with organoid characterisation and troubleshooting, Thomas Theil and Thomas Pratt for helpful comments on the manuscript. This work was supported by Simons Initiative for the Developing Brain (grant number 529085, JOM and DJP) and the Medical Research Council (grant number MR/Y004299/1, WKC, JOM and DJP).

---

**Supplementary Fig 1.**
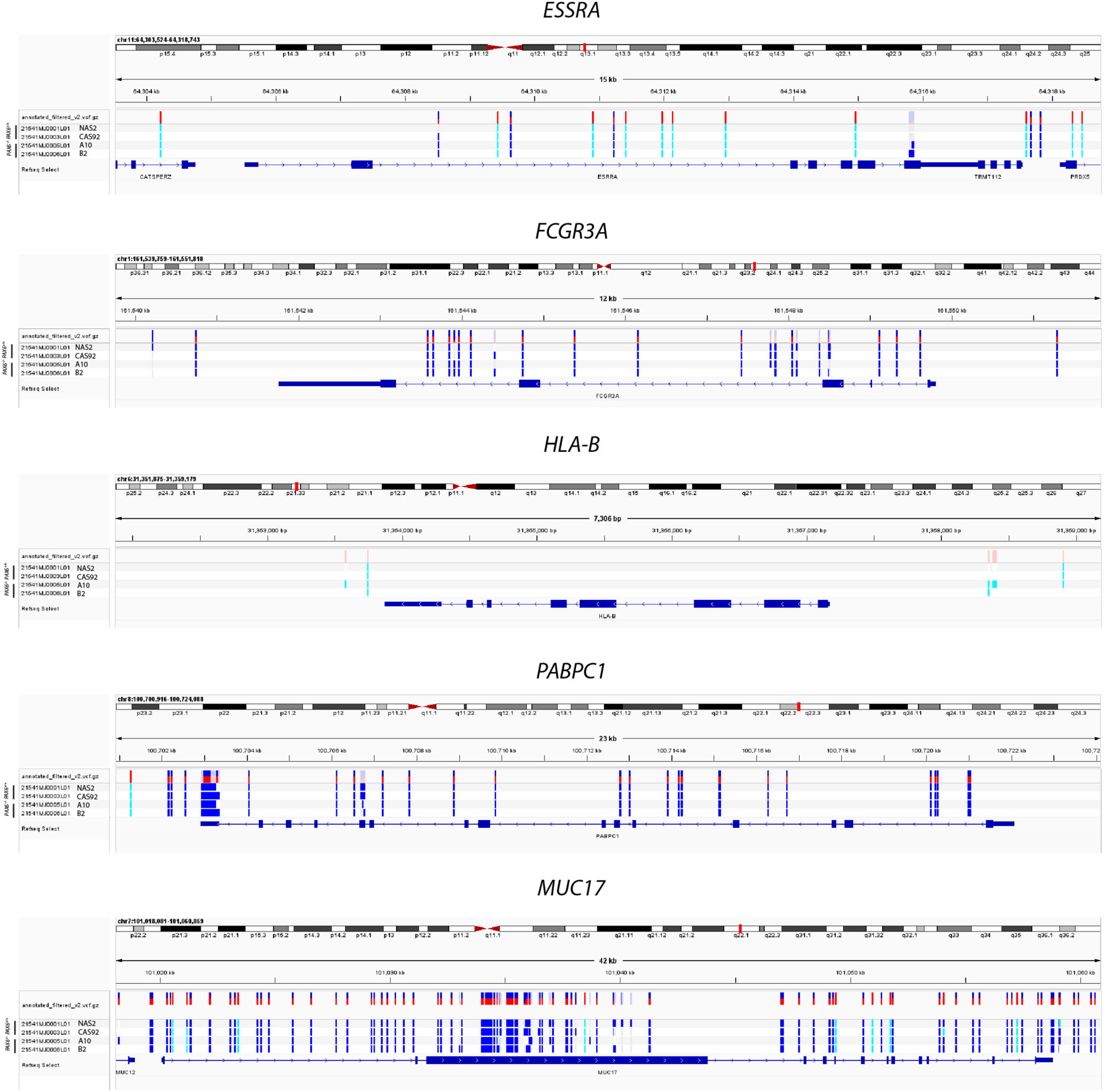
Integrated Genomics Viewer (IGV) plots of genes with the highest difference in mutation counts (SNV, indels, and point mutations) between *PAX6^+/+^* and *PAX6^-/-^* groups. Mutation variants for each individual samples are denoted as: homozygous variant – cyan; heterozygous variant – blue while allele frequencies are denoted as: high allele frequency of the variant – red; high allele frequency of reference sequence – blue. We examined five genes that showed the highest difference in mutation counts between *PAX6^+/+^* and *PAX6^-/-^* which were *MUC17, PABPC1, ESSRA*, *FCGR3A* and *HLA-B*. *MUC17* mutations are commonly found in iPSCs and have no reported effects on pluripotency or genomic integrity (D’Antonio et al., 2018). Other than *MUC17*, no mutations were found in coding regions of any of the other genes. Moreover, the variants reported by WGS were found in all four cell lines (Supplementary Fig 1) signifying that they are background mutations <u>(Young et al., 2012)</u> present in the parental line NAS2.

**Supplementary Fig 2.**
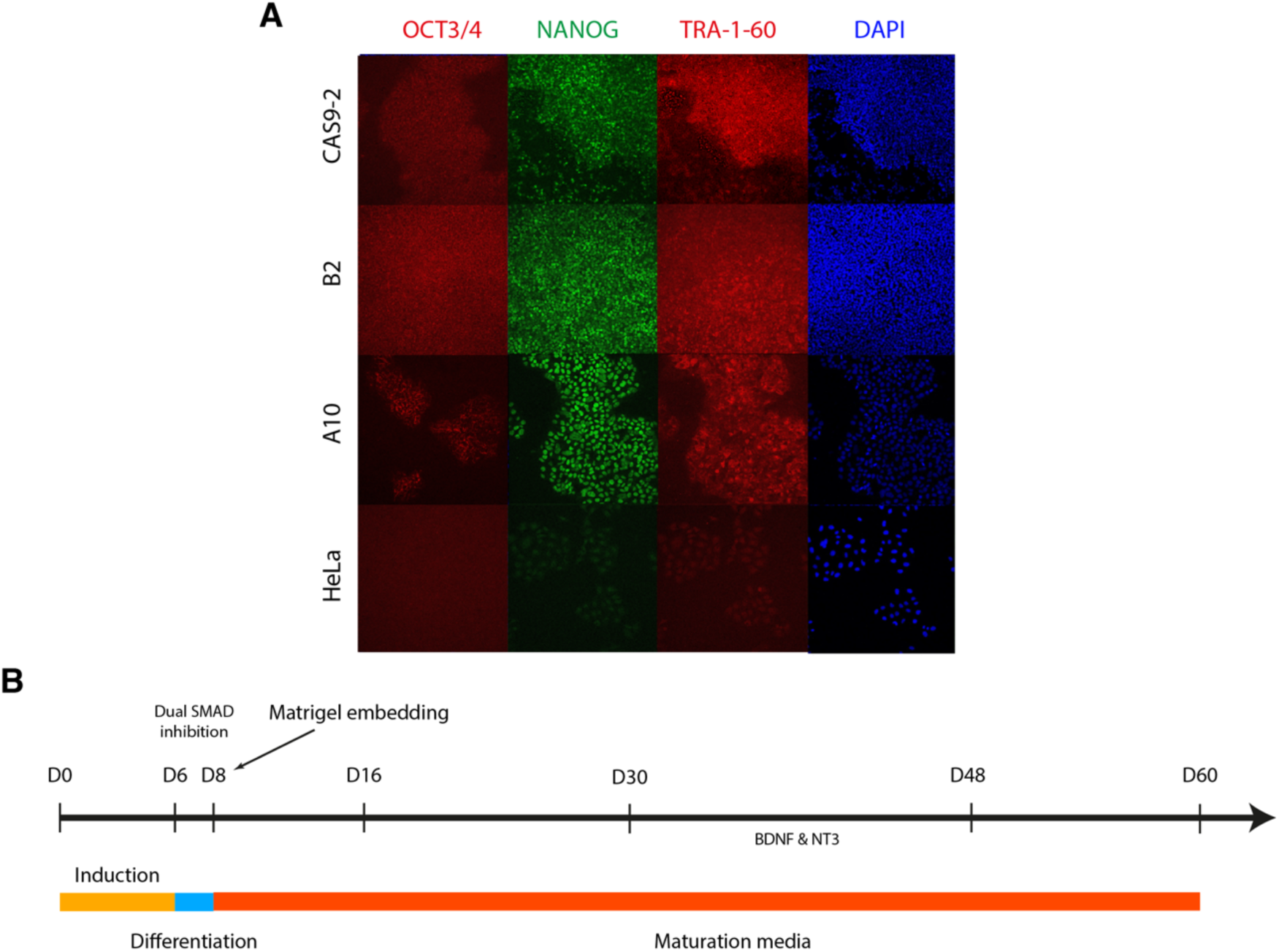
(A) Immunostaining of CRISPR edited iPSC cell lines used in this study showing expression of pluripotency markers NANOG, OCT3/4 and TRA-1-60. HeLa cells were used as a negative control. (B) Timeline illustrating organoid differentiation protocol.

**Supplementary Fig 3.**
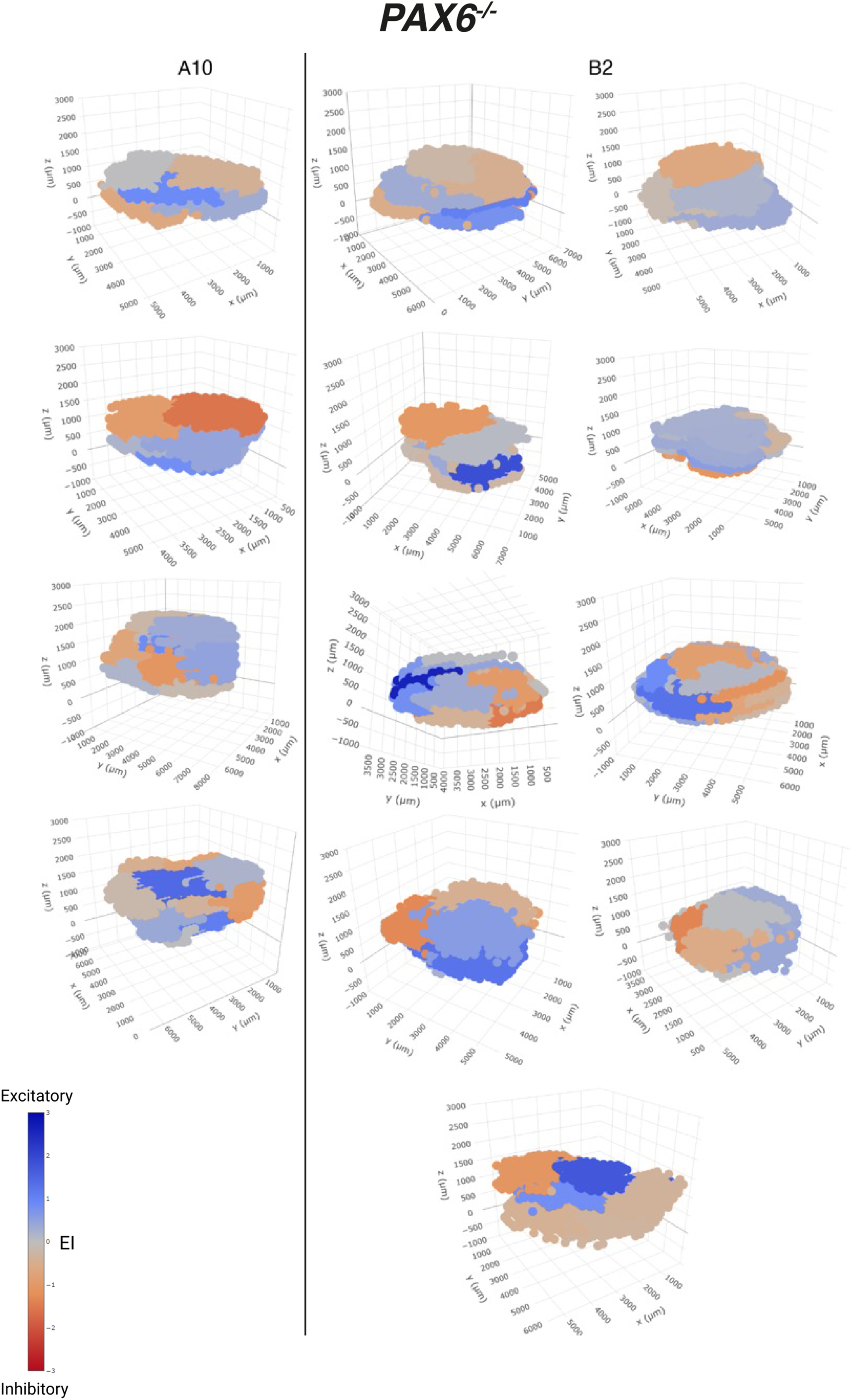
3D models of all individual organoids of both *PAX6^-/-^* clones (A10 and B2) that were quantified.

**Supplementary Fig 4.**
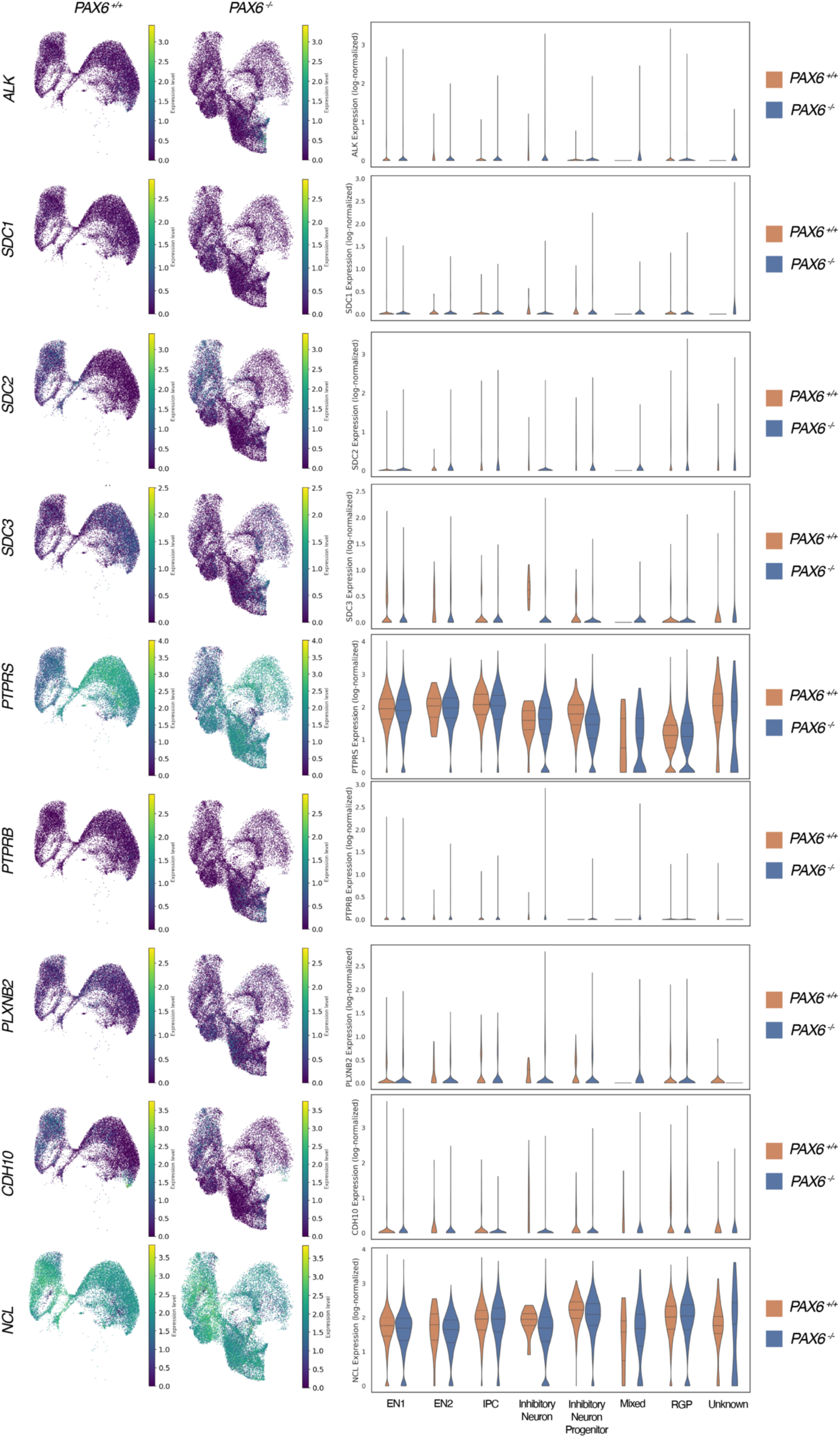
UMAP and violin plots showing expression of genes involved in the PTN signalling pathway showing no differences between *PAX6^+/+^* and *PAX6^-/-^*.

**Supplementary Fig 5.**
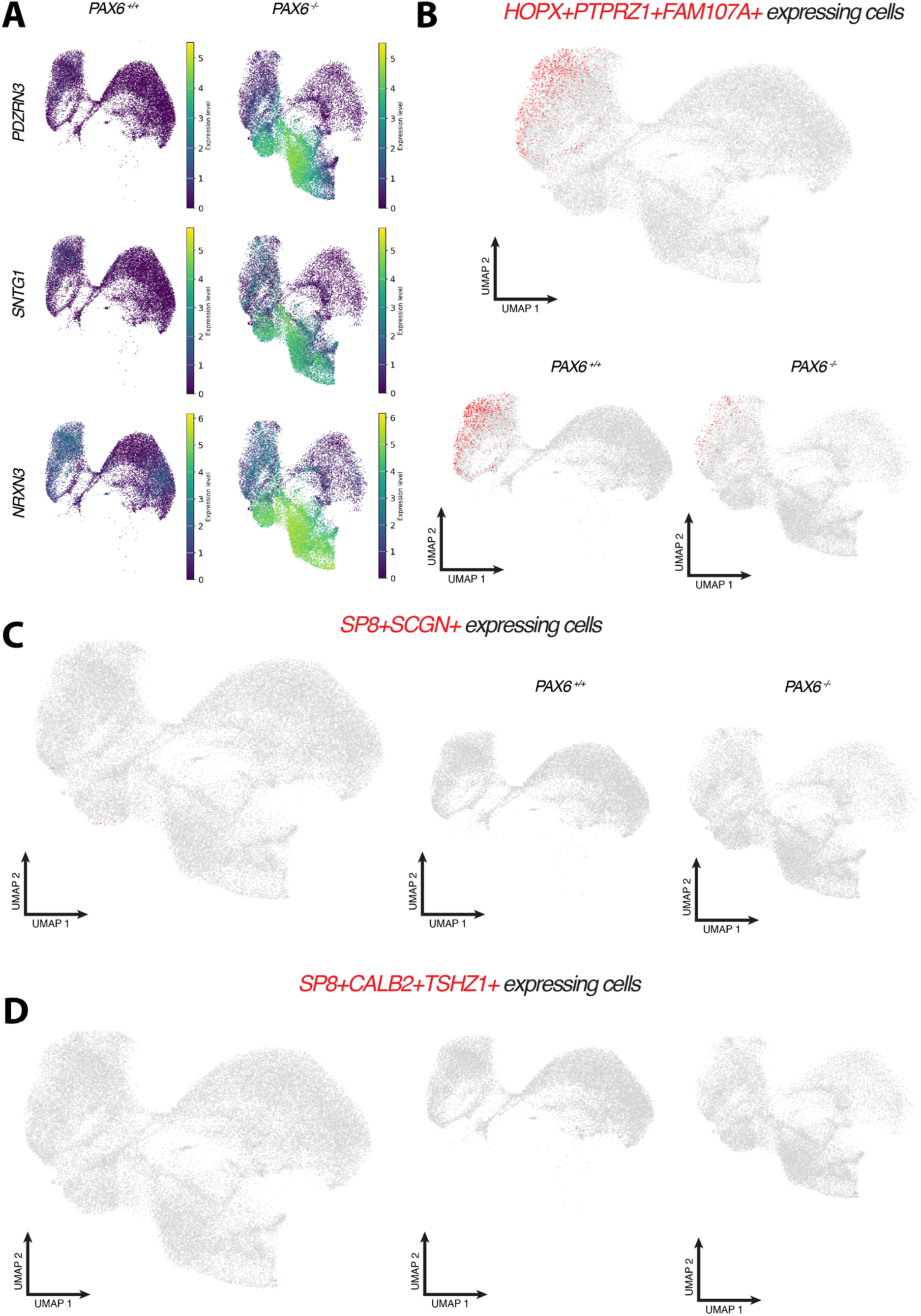
(A) UMAP plots of cells expressing *PDZRN3*, *SNTG1* and *NRXN3* with higher expression in yellow and low expression in dark blue. (B) UMAP plots showing cells that express oRG markers *HOPX*, *PTPRZ1*, and *FAM107A* (C) UMAP plots showing cells that co- express *PAX6* and *MEIS2* (D) UMAP plots showing cells that co-express *SP8* and *SCGN* (E) UMAP plots showing cells that co-express *SP8*, *CALB2* and *TSHZ1*.

**Supplementary Table 1.**
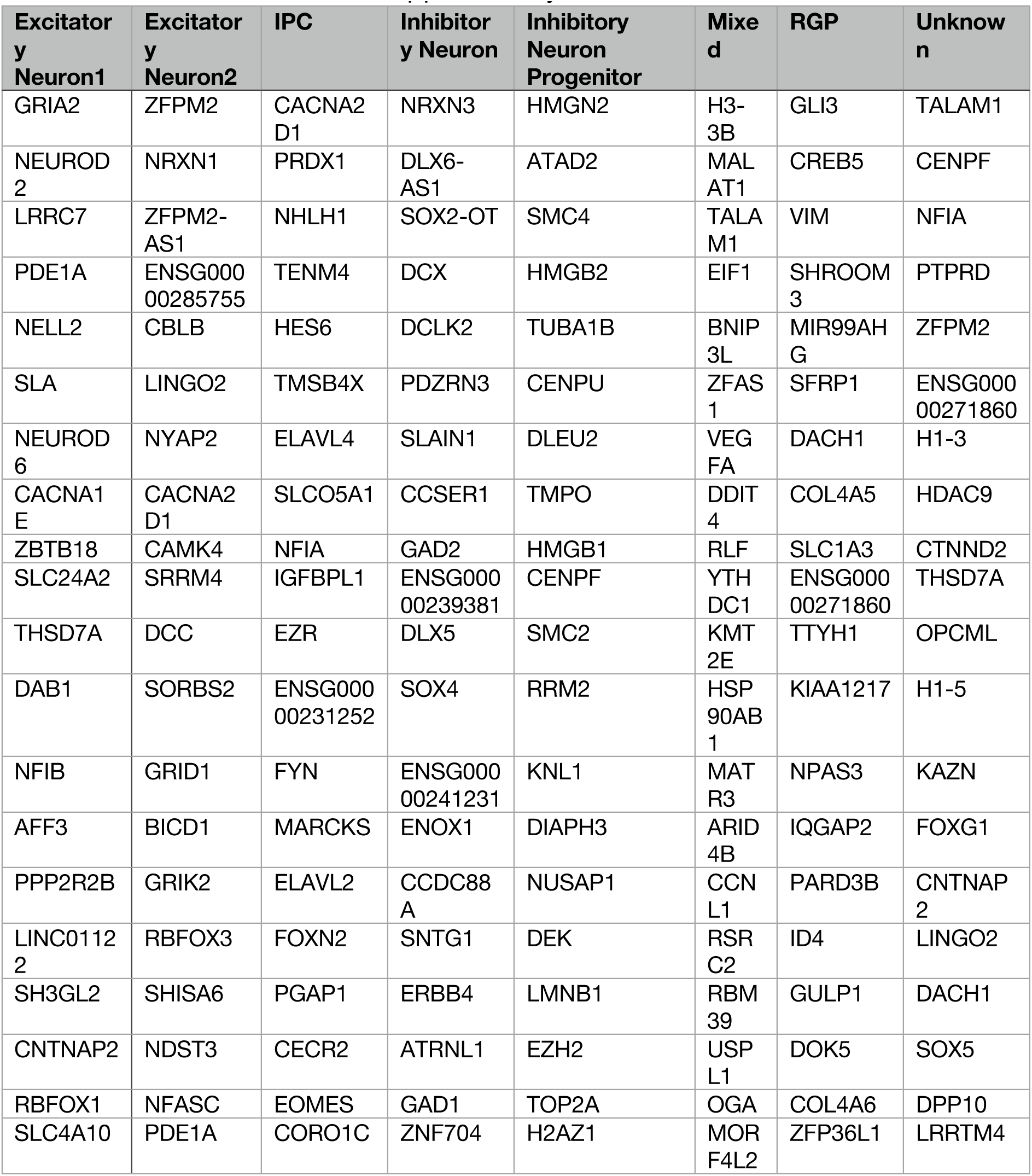
Marker genes of each cell type.

